# Computational principles of neural adaptation for binaural signal integration

**DOI:** 10.1101/863258

**Authors:** Timo Oess, Marc O. Ernst, Heiko Neumann

## Abstract

Adaptation to statistics of sensory inputs is an essential ability of neural systems and extends their effective operational range. Having a broad operational range facilitates to react to sensory inputs of different granularities, thus is a crucial factor for survival. The computation of auditory cues for spatial localization of sound sources, particularly the interaural level difference (ILD), has long been considered as a static process. Novel findings suggest that this process of ipsi- and contra-lateral signal integration is highly adaptive and depends strongly on recent stimulus statistics. Here, adaptation aids the encoding of auditory perceptual space of various granularities. To investigate the mechanism of auditory adaptation in binaural signal integration in detail, we developed a neural model architecture for simulating functions of lateral superior olive (LSO) and medial nucleus of the trapezoid body (MNTB) composed of single compartment conductance-based neurons. Neurons in the MNTB serve as an intermediate relay population. Their signal is integrated by the LSO population on a circuit level to represent excitatory and inhibitory interactions of input signals. The circuit incorporates an adaptation mechanism operating at the synaptic level based on local inhibitory feedback signals. The model’s predictive power is demonstrated in various simulations replicating physiological data. Incorporating the innovative adaptation mechanism facilitates a shift in neural responses towards the most effective stimulus range based on recent stimulus history. The model demonstrates that a single LSO neuron quickly adapts to these stimulus statistics and, thus, can encode an extended range of ILDs in the ipsilateral hemisphere. Most significantly, we provide a unique measurement of the adaptation efficacy of LSO neurons. Prerequisite of normal function is an accurate interaction of inhibitory and excitatory signals, a precise encoding of time and a well-tuned local feedback circuit. We suggest that the mechanisms of temporal competitive-cooperative interaction and the local feedback mechanism jointly sensitize the circuit to enable a response shift towards contra-lateral and ipsi-lateral stimuli, respectively.

**Author summary:** Why are we more precise in localizing a sound after hearing it several times? Adaptation to the statistics of a stimulus plays a crucial role in this.

The present article investigates the abilities of a neural adaptation mechanism for improved localization skills based on a neural network model.

Adaptation to stimulus statistics is very prominent in sensory systems of animals and allows them to respond to a wide range of stimuli, thus is a crucial factor for survival. For example, humans are able to navigate under suddenly changing illumination conditions (driving a car into and out of a tunnel). This is possible by courtesy of adaptation abilities of our sensory organs and pathways.

Certainly, adaptation is not confined to a single sense like vision but also affects other senses like audition. Especially the perception of sound source location. Compared to vision, the localization of a sound source in the horizontal plane is a rather complicated task since the location cannot be read out from the receptor surface but needs to be computed. This requires the underlying neural system to calculate differences of the intensity between the two ears which provide a distinct cue for the location of a sound source. Here, adaptation to this cue allows to focus on a specific part of auditory space and thereby facilitates improved localisation abilities.

Based on recent findings that suggest that the intensity difference computation is a flexible process with distinct adaptation mechanisms, we developed a neural model that computes the intensity difference to two incoming sound signals. The model comprises a novel mechanism for adaptation to sound source locations and provides a means to investigate underlying neural principles of adaptation and compare their effectivenesses. We demonstrate that due this mechanism the perceptual range is extended and a finer resolution of auditory space is obtained. Results explain the neural basis for adaptation and indicate that the interplay between different adaptation mechanisms facilitate highly precise sound source localization in a wide range of locations.

## Introduction

Localization of a sound source in space is a crucial task for every living being that is facilitated with an auditory system, may it be for hunting or predator avoidance. In contrast to visual object localization which is limited to a certain range for most animals, auditory localization allows for detection of sound sources in 360°. It works in low contrast environments (bright day light or complete darkness) and some animals solely rely on auditory localization to hunt and find their way through an environment [1, 2]. Humans take advantage of their accurate sound source localization system e.g. to focus on a single speaker among plenty other sources (see *cocktail party effect* [3]).

Auditory localization is a rather complicated task, since a sound signal does not explicitly convey information about its location. Instead, lateral localization predominantly relies on two binaural cues: the interaural time difference (ITD) and the interaural level difference (ILD). The ITD is based on the delay of signals between the two ears. That is, for sound source locations on the ipsilateral side of the head sound waves are delayed at the contralateral ear due to the longer travel distance, thereby creating a difference in the arrival time of the signal between the two ears. Such difference can be computed by a neural coincidence mechanism [4]. However, more recent findings suggest that in mammals the encoding of such time differences is achieved by a complex inhibition circuit [5, 6]. The ILD, on the other hand, exploits the fact that high frequency sounds coming from the contralateral side are attenuated by the head and thereby are reduced in their intensity on the ipsilateral side [7]. The amplitude difference is computed by comparing the different intensity levels of the left and right ear.

The superior olivary complex, a midbrain region that connects the cochlear nucleus (CN) and the inferior colliculus (IC), is the first stage in the auditory pathway that receives inputs from the ipsilateral and contralateral ear and thereby is the first region of binaural processing. Particularly the lateral side of the superior olive (LSO) receives excitatory input form the ipsilateral CN and inhibitory input from the ipsilateral medial nucleus of trapezoid body (MNTB), a relay station for signals coming from the CN of the contralateral side. That is, for a high intensity signal at the ipsilateral ear, LSO neurons are strongly excited. However, if the intensity of the signal at contralateral ear becomes stronger, inhibition on LSO neurons is increased and their activity reduced. By integrating these signals neurons in the LSO compute the ILD between the two ears [8–11].

From the stiffness change of hair cells in the cochlea to noise suppression in the auditory cortex, the auditory system implements several adaptation mechanisms throughout its pathway. Recent investigations indicate that also the process of binaural signal integration in the LSO is highly dynamic and facilitated by a dedicated mechanism that allows it to rapidly adapt to statistics of perceived stimuli. Thereby providing a dynamic encoding of perceptual auditory space, which allows to maintain high sensitivity to sound source locations [12–15].

Here, we investigate adaptation ability of LSO model neurons to perceived stimuli statistics. These neurons exhibit two distinct adaptation mechanisms: an implicit and explicit one. The former emerges from the sensitivity of LSO neurons to differences of signal arrival time [16, 17]. The physiological nature of this mechanisms allows it to work instantly. The latter one is realized by a retrograde GABA signaling pathway [18] and works in the magnitude of seconds.

By adapting to recently perceived stimuli, the (neural) system potentially either increases its population response efficiency [19] or improves its localization precision for consecutive stimuli, since it is likely that two consecutive sounds originate from nearby spatial locations. In other words, adapting to a previous stimulus can increase localization ability for the consecutive one. This behavior was shown for human participants in a behavioral experiment in which they had to localize sound sources after being exposed to an adapter tone [12, 20]. In addition, the authors conducted neurophysiological experiments with gerbils that led them to the conclusion that this adaptation mechanism is mediated by a “retrograde GABA signaling pathway” that inhibits the afferents of an LSO neuron and thereby adapts the neuron’s responses based on its recent activity.

How these adaptation mechanisms could be realized in a neural network model and to what extend they support and even improve the localization of sound sources is investigated in the following. The implemented neural model serves as an example on how such a mechanism can realize a canonical framework of synaptic adaptation with the potential to take place in other tasks where adaptation is required. Simulations indicate that this adaptation enables a large increase in the effective operational range while simultaneously providing higher resolution of the spatial location of interest in the perceptual auditory space.

We develop a computational model of the neural dynamics for adaptive shift estimation in binaural sound localization which supposedly takes place in the LSO. The model neurons utilize conductance-based properties of state change from excitatory and inhibitory inputs. Fast adaptation is achieved by a local control mechanism based on an activity dependent GABA pathway which adapts the effective synaptic integration at the basal dendrites of LSO neurons.

A series of computational experiments are reported to investigate efficacies of various adaptation mechanisms of model neurons and to quantitatively analyze their functionalities.

## Model

In this section we introduce the architecture of the model which components and interactions are based on neurophysiological findings. The LSO is the first stage of binaural processing in the auditory pathway. It receives direct ascending inputs from spherical bushy cells (SBC) of the ipsilateral CN and indirect inputs from globular bushy cells (GBC) of the contralateral CN, via the MNTB. The MNTB serves as a relay station that converts the excitatory signal of GBC neurons to an inhibitory input to the LSO. In contrast, SBC neurons of the ipsilateral CN project directly to the LSO and provide excitatory input. By integrating these two signals, neurons of the LSO encode the ILD by means of a monotonic correspondence between response rate and ILD of the signal. Taken into account this physiological hierarchy the presented model consists of these two distinct neuron populations and their circuit-level interactions as shown in Fig. 1.

**Fig 1.**
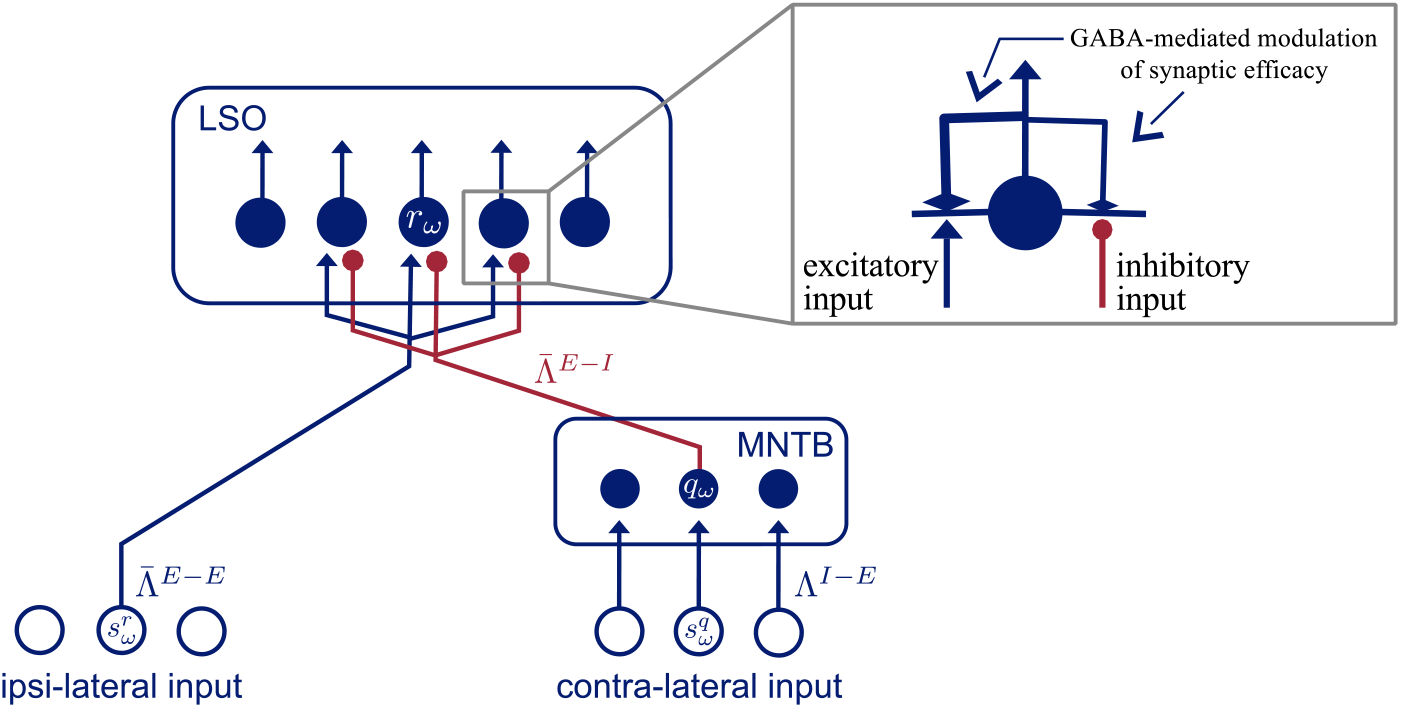
Architecture of model. Blue filled circles represent model neurons. Blue empty circles indicate inputs to model neurons. Red bullet-headed connections are inhibitory and blue arrow-headed connections are excitatory connections from inputs to neurons and from neurons to other neurons, respectively. Top right corner is a enlarged view on the synaptic connections of an LSO neuron. The output of such a neuron modulates its afferent connections.

Each of these populations comprises an array of *N* neurons, each selective to a specific frequency band centered at frequency *ω* (the neuron’s characteristic frequency). Their response is formally described by a first-order ordinary differential equation of the change of its membrane potential (see [21, 22]). The input conductances to a modeled LSO cell are defined by 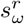 and 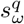. These inputs represent the activity of SBC and GBC cells of the ipsi- and contralateral CN, respectively, with characteristic frequency *ω* = {1, …, *N*}. Thereby inputs to the model as well as the neurons within a population are tonotopically ordered. The input signals to the model are assumed to emerge from neuron populations of preceding stages of the LSO in the auditory pathway, namely spherical and globular bushy cells in the ipsi- and contralateral cochlear nucleus, respectively. It has been shown that the activity of these cells is a linear function of the sound level in dB (logarithmic scale) of perceived stimuli [23]. For simplicity, it is thereby assumed that the inputs are in the range 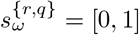 and that the values linearly correspond to the perceived level intensities.

### LSO neurons

The modeled LSO neuron population receives ipsi-lateral excitatory input from SBC cells and inhibitory input from neurons of the contra-lateral MNTB population. We employ a single-compartment model to describe the membrane potential, or activity, of LSO neurons *r_ω_* which are selective to frequency bands *ω*. We defined activities according to the leaky integrator equation to model membrane potential dynamics with the membrane current as the sum of excitatory, inhibitory, and leak conductances [24–26]. Similar neuron models have previously been demonstrated to successfully describe neuron and network characteristics in a variety of vision tasks, e.g., [27, 28]. The equation reads

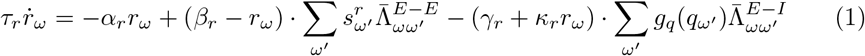

where 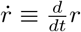 and *τ* define the membrane time constant of the neuron. The input current (right hand side) is defined by a leak current (first term) and lumped excitatory and inhibitory currents (second and third term), respectively. We arbitrarily set the passive equilibrium potential to zero with constant passive leak conductance *α_r_*, which leads to passive decay to zero resting level if inputs are switched off. In the second summation term, 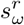 describes the ipsilateral excitatory input to an LSO neuron arising from a driving input population of excitatory SBC neurons which are integrated by a weighted summation with an interaction kernel 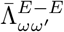 over frequencies (index *E − E* denotes excitatory-to-excitatory connections). The third summation term integrates activities of inhibitory MNTB neurons, with activity *q_ω_*, by a weighted summation with an interaction kernel 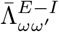 over frequencies (index *E* − *I* denotes inhibitory-to-excitatory connections). Inhibitory input *q_ω_* (see Eq. 6) is passed through a nonlinear activation function *g_q_*(•). Together, excitatory and inhibitory conductances realize a recurrent on-center/off-surround competitive mechanism in a dynamic field of neurons [26]. The driving forces in the excitatory and inhibitory input currents are defined by the excitatory saturation level *β_r_* and the inhibitory saturation level *γ_r_/κ_r_*, respectively. The latter constants determine the strength of the subtractive and the divisive influence of the inhibitory input current.

The key feature of the model is a retrograde GABA signaling circuit [18] that attenuates the efficacy of the model’s afferent connections, where the individual effectiveness depends on the activity of a single LSO neuron. This adaptive inhibition is asymmetric exerting stronger inhibition on excitatory synaptic input than on inhibitory synaptic input. To account for these effects, we incorporated an activity-dependent modulation function that decreases the effective weight coefficients. Formally, we replace the weight kernels 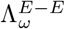 and 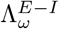 by state-dependent adaptive terms

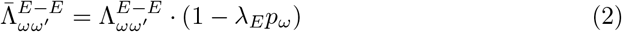

and

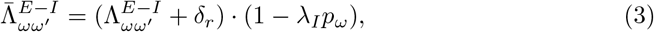

respectively. The parameter *δ_r_* denotes an offset level for inhibitory GABA signaling strength and *λ_E_* and *λ_I_* are the GABA receptor effectivenesses on the excitatory and inhibitory input, respectively. We will discuss these parameters and their origin in more detail in section GABA parameter strength. The state variable *p_ω_* models the GABA receptor activation over time and can be described by

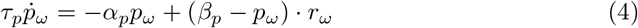

where parameter *τ_p_* defines the membrane capacity, *α_p_* is a decay rate and *β_p_* describes a saturation level of the inputs. The net effect of this weight adaptation in the kernels is a monotonic reduction of the efficacy of the excitatory and inhibitory inputs of an LSO neuron. The down-modulatory effect on the effective weights is controlled by the activity of the LSO neuron itself. This mechanism allows the neuron to adapt to stimulus history and thereby provide a means for dynamic adaptation.

The firing rate of an LSO neuron is calculated by a sigmoidal activation function *g_r_* of its membrane potential *r_ω_*

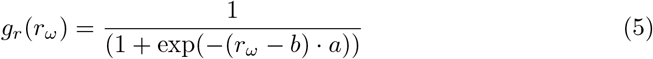

### MNTB neurons

Model MNTB neurons receive excitatory input from GBC of the contralateral CN and project an inhibitory signal to the ipsilateral LSO [29]. Thereby, MNTB neurons are a relay station between the ipsi- and contralateral hemisphere. The inhibitory input *q_ω_* from relay cells in the MNTB to the LSO is defined by

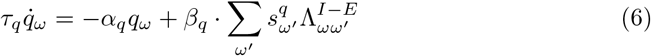

where 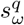 describes the contralateral input to an MNTB neuron, arising from an input population of GBC neurons and weighted by interaction kernel 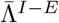. Parameters *τ_q_*, *α_q_* and *β_q_* have same functionality as in Eq. 1. The interaction kernels of the neuron inputs in Eq. 1 and 6 are normalized Gaussians with different sigmas *σ^i^*, *i* ∈ {*E* − *E*, *E* − *I*, *I* − *E*} depending on the given connection (see table 1).

The firing rate of an MNTB neuron is defined by the rectified signal of its membrane potential

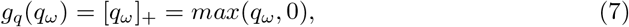

where *max*(*q_ω_*, 0) denotes the half-wave rectification of the activation.

**Table 1.** Model parameters

| General Parameters |  |  |  |
| --- | --- | --- | --- |
| N (# Neurons) | 5 |  |  |
| sigmoid parameter $a$ | 20 | | |
| sigmoid parameter $b$ | 0.2 | | |
| LSO Neuron |  |  |  |
| $\tau_r$ | 0,025 | $\lambda_I$ | 1.0 |
| $\alpha_r$ | 1.0 | $\lambda_E$ | 2.0 |
| $\beta_r$ | 1.0 | $\sigma^{E-E}$ | 0.5 |
| $\gamma_r$ | 3.0 | $\sigma^{E-I}$ | 0.6 |
| $\kappa_r$ | 4.0 | $\delta_r$ | 0.16 |
| MNTB Neuron |  |  |  |
| $\tau_q$ | 0,025 | $\sigma^{I-E}$ | 0.1 |
| $\alpha_q$ | 2.0 | | |
| $\beta_q$ | 1.0 | | |
| GABA Receptor |  |  |  |
| $\gamma_p$ | 2500.0 | | |
| $\alpha_p$ | 25.0 | | |
| $\beta_p$ | 125.0 | | |

## Results

Model simulations in this section provide a basis for evaluating binaural stimulus integration and a quantitative analysis of the adaptation efficacy of model neurons simulating LSO functions.

In experiment 1 general response properties of the model under different neuron parameter conditions are presented. We show that the response range and the coding precision of model neurons can be adjusted by altering parameter values that control how inhibitory inputs are integrated to calculate the membrane potential. Having demonstrated that, we introduce a retrograde GABA signaling circuit to the model that dynamically alters the effective strength of the inhibitory inputs based on recent neuron activity.

Thus, the experiment 2 inspects the response adaptation for this local GABA inhibition circuit. The strength of the inhibitory GABA signals imposed on the excitatory and inhibitory inputs determines the efficacy and shape of the adaptation. By systematically varying parameters that control the strength of the inhibitory GABA signals, an optimal value for the ratio of the parameters is identified on the basis of a novel adaptation index. This adaptation index (similar to the *standard separation* in [30] and [20]) quantifies the coding precision of a neuron with respect to its ability to register relative (azimuthal) offsets of sound sources, thereby creating a unique measure of the adaptation quality. This index is the basis for evaluation and comparison of the different adaptation processes and efficacies in the following experiments.

The effect of the introduced adaptation mechanism on response characteristics based on stimulus history is examined in experiment 3. Here, the influence of stimulus history on location estimation is measured by presenting adapter tones with fixed ILDs shortly followed by test stimuli with different ILDs, similar to behavioral experiments in [20] and [12].

Experiment 4 characterizes a second adaptation process in LSO neurons: the influence of time delays between the ipsilateral and contralateral inputs on the adaptation capacity of the model neuron (time-intensity trading [17, 31]). It demonstrates that the offset of arrival time between the input signals plays a crucial role in the integration process and facilitates higher localization accuracy.

All in the following presented results are calculated from the network responses readout at a single neuron level after keeping the input stimuli constant for at least 400 time steps. This duration is sufficient for the neuron to dynamically converge to its equilibrium membrane potential of numerical integration of the state equations (see Fig. 2**B** inset).

**Fig 2.**
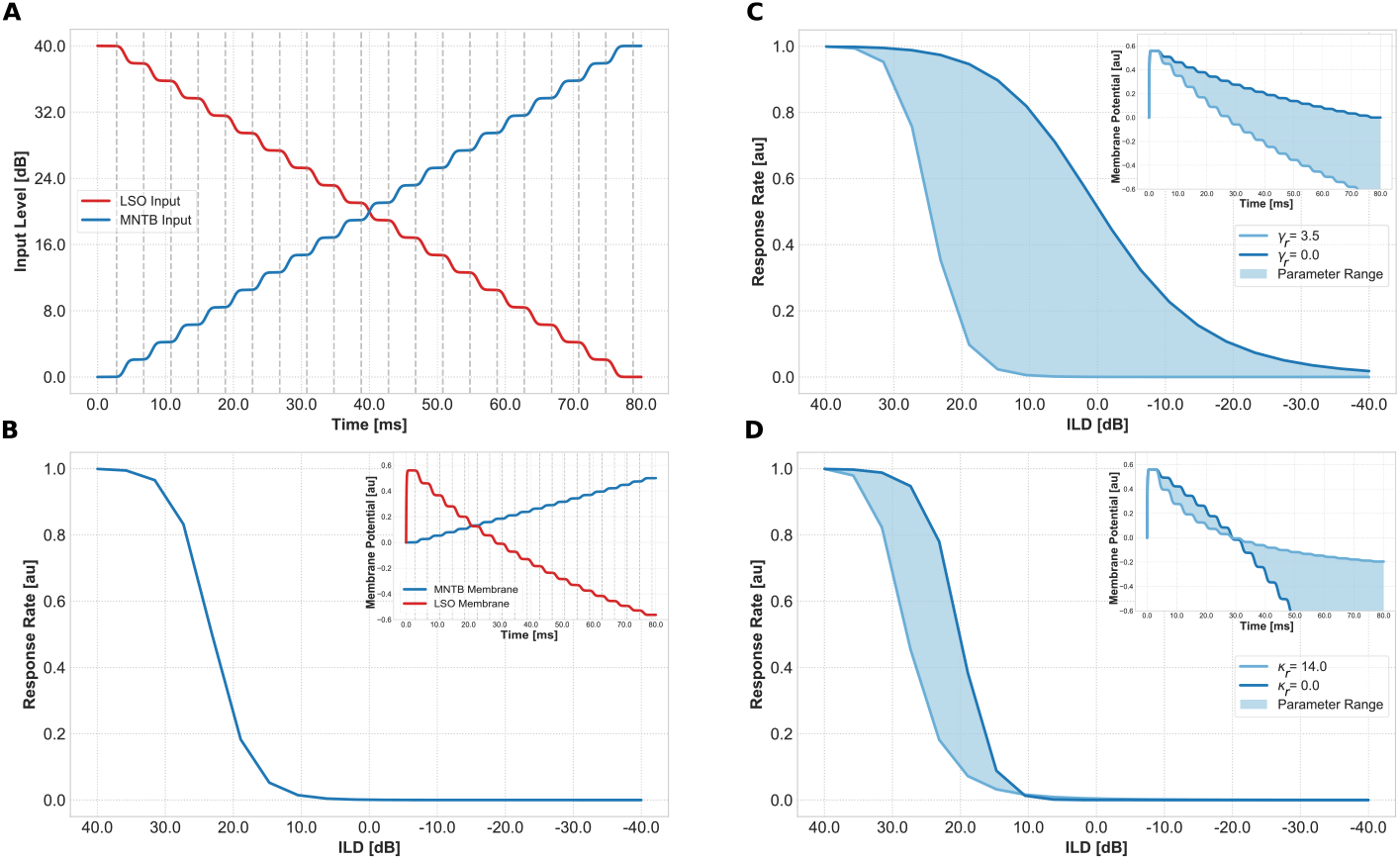
Input generation and parameter influences. (A) Input generation for the model system for ILD calculation. Red line shows excitatory input from cells of the ipsilateral cochlear nucleus to LSO neurons. Blue line shows excitatory input from cells of the contralateral cochlear nucleus to MNTB neurons which are converted to inhibitory input to LSO neurons. Vertical grey dashed lines indicate measurement points of neuron’s response. (B) Model response of an LSO neuron for a default set of parameter values. Blue line shows the response rate of a single LSO neuron to stimuli as presented in (A) over interaural level difference. The ILD is determined by subtracting the level of the ipsilateral stimulus from the level of the contralateral stimulus. Inset shows membrane potentials for LSO (activity *r*, red) and MNTB (activity *q*, blue) neurons. Probing times (grey dashed lines) are chosen to ensure that both membrane potentials have reached a steady state before measuring. These states can be identified by horizontal plateaus in the display where the membrane potential is constant. (C,D) Model response for parameter values *γ_r_* and *κ_r_*, respectively. (C) Parameter values of *γ_r_* are varied in range [0.0, 3.5] which control the strength of subtractive inhibition. (D) Model responses for different *κ_r_* values in range [0.0, 14] that regulates the effect of shunting inhibition on the membrane potential. Filled areas are approximated from simulation results.

In particular, a constant level is presented for 400 time steps to the ipsilateral and contralateral input populations before changing it. This presentation time corresponds to the reaction time of neurons and is consistent throughout all experiments expect experiment number 4. For the numerical integration of the state equations we chose *Euler’s method* with a step size of ∆*t* = 0.001 (for details see [32]).

The choice of model parameters of the neurons is crucial for their response behavior in terms of the effective response range (steepest slope) and of the range of dynamic adaptation. The parameters for the following simulations are chosen to fit a variety of neurophysiological experiments. In particular, we focused on the results of the dynamic adaptation experiment [18] (experiment 2 and 3), where the authors demonstrate that activation of a retrograde GABA signaling pathway leads to a shift of LSO neurons’ response curves, and the input timing experiment [16] (experiment no. 4), in which it is shown that LSO neurons respond to the ITD of the presented signals thus exhibiting time-intensity trading.

### Parameter influence and response behavior

Neural models have several parameters (e.g. membrane saturation, decay rate) that can be tuned to achieve a desired response behavior of a model neuron. Since we are investigating the integration of two incoming signals, the most influential parameters are the ones that control the effective strength of those signals on the membrane potential.

Therefore, the first experiment investigates the influence of model parameter values that control the influence and strength of inhibitory inputs. We demonstrate that the typical response curve of LSO neurons (sigmoidal function over ILDs) is shifted and that its slope is modulated by these parameters. In addition, the experiment shows the *default response* of a model LSO neuron that is used for comparison in further experiments. Since the purpose of this experiment is to investigate influence of model parameters without the influence of dynamic adaptation, GABA receptor parameters *λ_E_*, *λ_I_* and *δ_r_* are set to 0, effectively disabling this retrograde GABA signaling circuit (see Eq. 2 and Eq. 3).

Input stimuli to the neuron model over time for an example frequency band *ω* are shown in Fig. 2**A**. Note that all frequency bands are excited equally. Ipsi- and contralateral inputs are presented as monotonic functions of the sound level over time (decreasing for ipsilateral input and increasing for contralateral input, respectively). These input functions have several plateaus i.e. constant levels over time that allows the neuron’s membrane potential to reach a steady state (see Fig. 2**B** inset). Note, that the response of a single LSO neuron is the result of a dynamic process of direct excitatory inputs and indirect inhibitory inputs from MNTB neurons.

The output response of a model LSO neuron is computed by applying a sigmoid transfer function (see Eq. 5) to the membrane potential to generate a firing rate. The transfer function is tuned to have typical response properties of measured neurons in mammals [16, 33]. The observed output starts to increase for ILD values around +10*dB* and has its steepest slope for ILD values of 25*dB* before it saturates for a maximal perceivable ILD value (in this model +40*dB*). The response of the model for the default set of parameter values, as defined in table 1, is shown in Fig. 2**B**.

Since the maximal numerical input to our model neuron is scaled to be positive and lower than 1.0, the range of input level difference is [−1.0, 1.0] (i.e. an input value of 1.0 on the ipsilateral and an input value of 0.0 on the contralateral side and vice versa). We defined the range of perceivable ILD to be between 40*dB* and −40*dB* to be in line with physiological ILD values [17, 34].

Two major mechanisms of gain control in neuronal circuits exist: *shunting* and *subtractive inhibition*. The term shunting refers to its effect of locally reducing the input resistance and increasing its conductances, thereby reducing the amplitude of excitatory postsynaptic potentials (EPSP) [35]. Subtractive inhibition is achieved by hyperpolarization of the neuron’s membrane due to the opening of voltage gated potassium channels.

Figure 2**C** and 2**D** display the influence of the LSO membrane equation parameters *γ_r_* (upper plot) and *κ_r_* (lower plot) (Eq. 1) that define the effect and type of the inhibitory input on the membrane potential. In the present model the parameter *κ_r_* controls the strength of the shunting term. The algebraic solution of the steady-state response of this inhibition type results in divisive inhibition. For *γ_r_* = 0*, κ_r_* > 0 the inhibition becomes a pure shunting inhibition. The strength of subtractive inhibition can be controlled by the parameter *γ_r_* and for *γ_r_* > 0*, κ_r_* = 0 the inhibitory input operates in a pure subtractive manner. When *γ_r_* > 0*, κ_r_* > 0 the inhibitory input is a combination of both inhibition mechanisms (see [22] for more details on parameter influence).

The displayed responses are attained by keeping one of the parameters at its default value (*γ_r_* = 3 and *κ_r_* = 4) while varying the other. Increasing the *γ_r_* parameter simultaneously leads to a steeper slope and a shift of the point of complete inhibition towards positive ILD values (louder tone at ipsilateral ear, see Fig. 2**C**). For different *κ_r_* values the slope of the response function is similar over all presented values (see Fig. 2**D**). Also the point of zero response stays constant. The only noticeable change is in the effective operating point, i.e. the point at which the neuron is maximally sensitive to changes. For small *κ_r_* values this point is closer to 20*dB* and shifts towards more positive ILDs for larger values. Since response to ILD values is highly variable among LSO neurons we fixed the values of the parameters to resemble neurophysiological response characteristics of an LSO neuron. That is, a maximal response close to the maximal perceivable difference and a point of zero response close to 10*dB* with a maximal sensitivity for sounds coming from the ipsilateral side.

The effect of the two parameters can also be seen in displays of the membrane potential (Fig. 2**C** and Fig. 2**D** insets), where parameter *κ_r_* defines whether the function decays in a convex or concave manner. The point of zero crossing does not change for different parameter values. In contrast, *γ_r_* changes the slope of the function (steeper slope for higher values) and by that shifts the point of zero crossing. The shape of the function however remains linear. For the influence of parameter values on coding precision see supplementary Fig. S1.

Nevertheless, there exist mechanisms and processes that can alter such a response curve of LSO neurons. One of these processes is mediated by GABAergic inhibition of the afferent connections. This mechanism facilitates a shift of the response curve to a great extent towards positive ILDs. We investigate this in the following experiment.

### Dynamic adaptation of response behavior

The following experiment two and three investigate the neural basis for dynamic adaptation in simulations of modeled LSO neurons. Recent neurophysiological findings revealed that neurons in the LSO employ a synaptic adaptation circuit that facilitates the neuron to adapt to recent stimulus history. This circuit has functional implications for sound localization as it shifts the range of highest sensitivity of the neuron towards recently perceived stimulus locations [12, 18, 20]. The underlying principle of this synaptic circuit is presumably the activation of pre-synatipc GABA receptors which regulate the effectiveness of excitatory and to a lesser extend inhibitory inputs. This mechanism depends on the activity of the LSO neuron itself and is in contrast to other gain control mechanism [36] not regulated by higher level (feedback) projections.

In their experiments with gerbils Magnusson and colleagues [18] applied the *GABA_b_* receptor agonist Baclofen during sound stimulation and measured the response rate of single LSO neurons. The application of Baclofen suppressed the response of the neuron compared to measurements before application of the drug (Fig. 1 in [18]). To inspect the location of such inhibition the investigators conducted further in vitro experiments with brain stem slices. These experiments confirmed that the inhibition takes place at the pre-synaptic connection by modulating the transmitter release. This leads to an attenuation of the connection strength and facilitates a gain control mechanism. Our model implements such a mechanism by means of an activity dependent reduction of excitatory and inhibitory input strength (see Eq. 2 and 3). The following experiments investigate the effect of the retrograde GABA signaling and its influence on stimulus history, by (i) systematically varying the effectiveness of the signal imposed at the excitatory and inhibitory input, respectively, and (ii) by measuring the influence of a preceding adapter tone on consecutive stimuli, similar to [20] and [12].

#### GABA parameter strength

In their study [18], the authors report that the response curve of an LSO neuron is shifted towards more positive ILD values for activated *GABA_b_* receptors (as shown in Fig. 3**B**). Two possible processes are conceivable that facilitate a shift of the response function towards positive ILD values: one is an increasing effect of the inhibitory input and therefore a greater suppression of the LSO neuron. However, *GABA_b_* is an inhibitory receptor and it is therefore unlikely to enhance inputs. The second, more likely, process suggests the suppression of the excitatory input which in turn leads to a greater influence of the inhibitory input. As a consequence, the response function is shifted towards more positive ILDs. The authors in [18] argued that the influence of the *GABA_b_* receptor activation at the excitatory and inhibitory inputs of the neuron differs and has a stronger effect at the excitatory projections.

**Fig 3.**
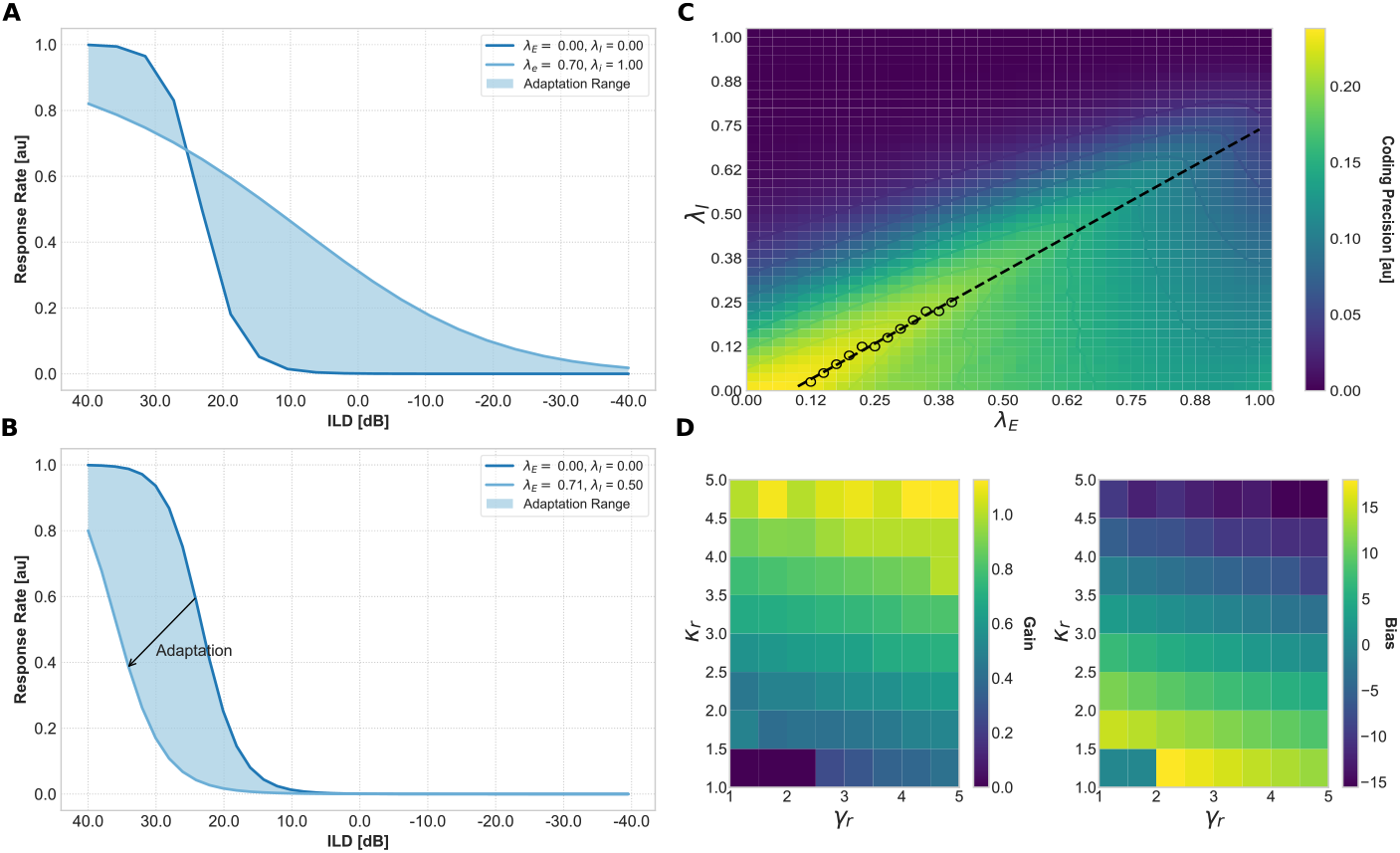
Effectiveness of GABA receptor on LSO response. (A) Adaptation range of model LSO neuron for inverse *GABA_b_* receptor effectiveness. Parameters *λ_E_* and *λ_I_* describe the inhibition effectiveness on excitatory and inhibitory inputs respectively. Normal (dark blue line) and shifted (light blue line) responses are shown. For *λ_E_* = *λ_I_* = 0 the ILD response curve is the same as the default response (Fig. 2**B**) and no dynamic adaptation process takes place. For *λ_E_* = 0.70 and *λ_I_* = 1.0 (inverse ratio) the response curve’s slope and consequently its sensitivity is decreased while the dynamic coding range is increased. (B) Adaptation range of model LSO neuron for increasing parameter values. Normal (dark blue line) and shifted (light blue line) responses are shown. Again, for *λ_E_* = *λ_I_* = 0 the ILD response function is similar to the default response. Light blue area indicates the adaptation range. (C) Exhaustive parameter evaluation of the *GABA* parameters *λ_E_, λ_I_*. Each point in the heat map indicates the coding precision value for a certain combination of *λ_E_* (abscissa) and *λ_I_* (ordinate). We assume that the neuron’s task is to achieve a preferably high coding precision value. Black dots depict these values (the maxima of the map). A linear function (dashed line) was fitted to those points and the slope (*m* = 0.80) and bias (*b* = −0.06) calculated. (D) Influence of model parameters *κ_r_* and *γ_r_* on the gain (left panel) and bias (right panel) of the linear regression.

To verify their argument, we investigate the response of a model LSO neuron for an inverse ratio of *GABA* receptor effectiveness (*λ_E_* < *λ_I_*). That is, we increase the effectiveness of *GABA* inhibition at the inhibitory synaptic inputs and reduce it at the excitatory synaptic inputs. This interaction leads to a more broadly tuned response function (see Fig. 3**A**). This is contradictory to neurophysiological findings where *GABA_b_* receptor activation leads to a finer tuning of the response curve. For our model we can conclude that in order to exhibit similar responses as measured in vitro, the effectiveness of *GABA* at the excitatory inputs needs to be stronger (*λ_E_* ≥ *λ_I_*). This is in line with results of in vitro experiments [18]. However, it is possible that the neural system takes advantage of this effect to increase the perceivable range of the LSO neuron (see Discussion for further details).

As shown in Fig. 3**A**, GABA effectiveness at the inhibitory input needs to be stronger to have the observed effect on LSO neurons (as presented in [18]). However, how much stronger the effect of GABA at the inhibitory input compared to the excitatory input needs to be to achieve an optimal adaptation to stimuli statistics is unknown.

Thus, experiment 2 explores the optimal ratio of the *GABA* inhibition efficacies on the inputs. This is achieved by an exhaustive parameter evaluation in which it is assumed that an optimal ratio entails maintaining a preferably high *coding precision* over the entire adaptation range. The *coding precision* is a measurement index adapted from the standard separation [37, 38] between two stimulus values and is defined as the derivative of the neuron’s response curve. Similar, the standard separation is defined by the difference of the firing rate of the two stimuli divided by the geometric mean of their standard deviations (see [19, 20] for details).

High coding precision values indicate steep slopes in the response curve of the neuron which, in turn, describe a high sensitivity of the neuron to changes in ILD values. Figure 3**C** depicts coding precision values for all parameter combinations *λ_E_*, *λ_I_* ∈ [0, 1]. Black dots indicate peaks in the coding precision for given *λ_E_* and *λ_I_* values and create a straight line with slope 0.80 and intercept −0.06 when linearly interpolated. These values are used as an indication for the GABA effectiveness on the inputs. We can conclude that the optimal ratio of *GABA* effectiveness at excitatory and inhibitory inputs in terms of coding precision is 1 : 1.25. Consequently, we choose model parameters in a way that the efficacy of *GABA* at the inhibitory input is 1.25 as high as for the excitatory input (*λ_E_* = *λ_I_* · 1.25) with an initial offset of *δ_r_* = −0.06.

Since values of model parameters *κ_r_* and *γ_r_* determine the ratio of *GABA* effectiveness, an exhaustive parameter sweep for different combinations of *κ_r_* and *γ_r_* (see supplementary Fig. S2) has been investigated. The gain and bias values for highest precision values are calculated for each combination (Fig. 3 **D**) as described above. For higher values of *γ_r_* the gain of the fitted line is increased whereas a change in *κ_r_* corresponds with a change in bias values.

The adaptation of the ILD response function for these parameters is depicted in Fig. 3**B**. For increasing parameter values the excitatory and inhibitory inputs are gradually inhibited which consequently shifts the ILD response function (blue area). The values *λ_E_* = 0.71 and *λ_I_* = 0.50 indicate the maximally effective inhibition since for higher values the response function did not shift further but remained constant. We take that as an indication for the maximal adaptation range facilitated by this gain control mechanism.

#### Neural adaptation to stimulus history

The third experiment tests the effect of the introduced gain control mechanism on localization quality. Previously conducted psychophysical experiments indicate that human subjects adapt to the history of recently presented stimuli [12, 20]. In a first study the human participants were tested with a 1*s* adapter tone of various ILD, consisting of broadband noise and an average binaural level of 60*dB*, followed by a 100 ms target tone of a constant ILD value over time [20]. The participants then had to indicate, in a two-alternative forced-choice task, whether the target sound was perceived on the left or right side of the midline (0*dB* ILD). Results show that the perceived midline of participants shifts in direction of the previously perceived adapter tone. The authors hypothesized that this is due to the effect that the brain attempts to maintain high sensitivity for locations of high stimulus density (here, the location of the previously perceived adapter tone). In addition, they report a response adaptation of neurons in the inferior colliculus (IC) of ferrets stimulated with a similar stimulus set as the human participants. Whether this adaptation process is constrained to IC neurons or takes place in a preceding processing stage, namely the LSO, was not investigated in this study. However, another study investigated neurons in the medial superior olive (MSO) for their adaptation capabilities to perceived sounds with different ITDs [39] and found similar adaptation capabilities.

Likewise, Lingner and colleagues [12] presented human participants with three different adapter tones (1*s* duration,0 μs, −181.4 μs or 634.9 μs ITD) followed by a target tone with different ITDs ranging from −634.9 to +634.9 μs. All presented sounds have a frequency of 200*Hz*. The authors reported a perceived shift of the target stimuli away from the adapter tone, indicating that the perceptual auditory space around the adapter tone is widened, leading to a higher sensitivity of sounds with ITDs similar to the adapter tone.

Taken together, the results of these studies demonstrate that the auditory system adapts dynamically to ILD and ITD on a behavioral level and comprises retrograde GABA signaling mechanisms in principal areas that process ITD and ILD information. However, whether LSO neurons exploit their retrograde GABA signaling mechanism to adapt to ILDs of previously perceived stimuli has not yet been shown. We hypothesize that this adaptation processes take places at the level of the LSO by retrograde signaling of GABA receptors as described in [18]. To test this hypothesis, the following experiment investigates the effect of stimulus statistics on model LSO neurons and their retrograde GABA signaling circuit. As in [20], an adapter tone is presented that is followed by a stimulus of various input level intensities. We expect that a preceding adapter tone activates the retrograde GABA circuit which attenuates the model’s response magnitude of the subsequent stimuli, thus shifting the response of the LSO neuron towards the ILD of the adapter tone. Note, that in contrast to the experiments in [20] we use the same stimulus type for the adapter tone as well as for the test tone since the sole purpose of the adapter tone is to activate the retrograde GABA signaling which is the case as long as the tone is within tuning characteristics of the receptive field (characteristic frequency and level difference) of the neuron.

The stimulus arrangement for this experiment is shown in Fig. 4**A**. It comprises an adapter tone of 1.2*s* with temporal ramps at the beginning and end of the sound followed by a 500*ms* period of silence. Subsequently, the target stimulus is presented with a duration of 2*s* and an ILD within the range as defined below. There are 41 adapter tones and 41 test stimuli of different level intensities. Hence, a total number of 41 × 41 = 1681 unique stimulus arrangements are presented to the model. The ILD of an adapter tone as well as a test stimulus ranges from −40*dB* to 40*dB*. Adapter tones are added and subtracted to the base line intensity of 20*dB*, on the contra- and ipsilateral side, thereby creating ILDs of −40*dB* and 40*dB*, respectively. For example, an adapter tone of 0 ILD indicates a sound level of 20*dB* on both sides whereas negative adapter tone levels decrease the input level on the contralateral and increase it on the ipsilateral side.

**Fig 4.**
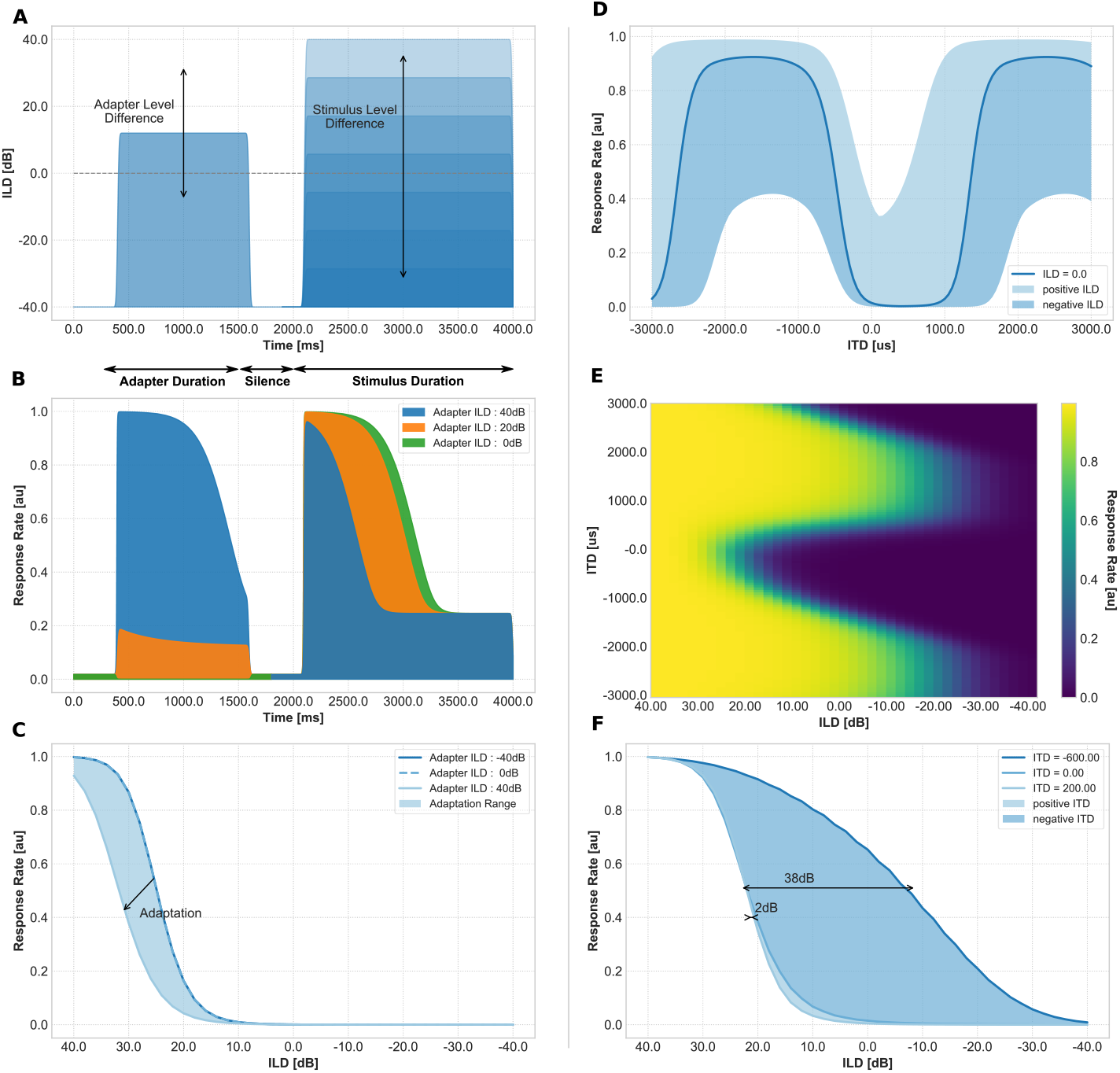
Adaptation of LSO neurons. Left column: stimuli and results of the adapter tone experiment 3. Right column: results of the timing experiment 4. (A) input stimulus example for Experiment 4. The first ridge depicts the ILD of the adapter tone with a duration of 1.2*s* followed by a break of 500*ms*. The second ridge shows the ILD of the actual test stimulus. Differently shaded areas illustrate different test stimulus ILDs for the same adapter tone. Grey dashed line marks 0*dB* ILD. (B) Responses of model neuron to different adapter tone intensities. Blue 40*dB*, orange 20*dB*, green 0*dB* and constant test stimulus intensity (40*dB*). The response of the neuron to an adapter tone sound level of 40*dB* (blue line) is maximal. However, the responses to the consecutive test stimuli is reduced through adaptation i.e. the higher the ILD of the adapter tone the lower the response to the test stimulus. (C) Response of a single LSO neuron to stimuli of constant ILD after the presentation of various adapter tone ILDs. Light blue dashed line depicts the response to a test stimulus of ILD 40*dB* after presenting an adapter ILD of 0*dB*. Dark blue line draws the response to the same test stimulus after presenting an adapter ILD of −40*dB*. No adaptation takes place and the response is similar to the default response of the neuron (compare Fig. 2**B**). Light blue line draws the attenuated response to the same test stimulus after presenting an adapter ILD of 40*dB*. (D) Responsiveness of LSO neuron for different ILDs over ITDs. The blue line indicates the neuron response for 0*dB* level difference, the darker blue area shows possible responses for higher ILD values, whereas the light blue area indicates possible responses for negative ILD values. The depicted ILD range is limited to −20*dB* and +20*dB* for a clearer plot. Note the u-shape (dark blue) and v-shape (light blue) response curves. (E) Neuron responses over various ITD and ILD values. (F) Adaptation range of model LSO neuron for various ITDs of the input signals. Light blue line and corresponding filled area indicates model response for stimuli with positive ITDs. Dark blue area shows responses for stimuli with negative ITDs. Area filled with blue is inferred from simulation results.

Responses of the LSO neuron model over time for three different adapter tone ILDs (40*dB,* 20*dB,* 0*dB*) and a constant test stimulus ILD of 40*dB* are shown in Fig. 4**B**. This test setup corresponds to a scenario in which the adapter tone source is varied between three different locations (approximately 90°, 45° and 0° azimuth) on the ipsilateral side and the test stimulus is located on the very far ipsilateral side (90° azimuth). The preceding adapter stimulus activates the retrograde GABA signaling circuit which subsequently attenuates the effectiveness of the neuron’s inputs. The response to the adapter tone stimulus accurately follows the input to the model. However, the response to the target stimulus (between 2000*ms* and 2500*ms*) is attenuated based on the ILD of the preceding adapter tone. As a consequence, this adaptation is leading to a shift of the model’s response curve towards the ILD of the adapter tone, as depicted in Fig. 4**C**. Measurements of the neuron activity are taken 100*ms* after stimulus onset to measure the initial response of the neuron. This elapse time was chosen because after the initial response the retrograde GABA circuit (*p_ω_*) is affected by the more recent activity of the neuron and the effect of the adapter tone vanishes. This is due to the time constant (*τ_p_*) of the adaptive term of the GABA circuit (see Eq. 4). For adapter tones with negative ILDs, i.e. located at the contralateral side, no adaptation takes place (the response is similar to the default response of the neuron) since the adapter tone’s ILD does not evoke a considerable activation of the GABA circuit of simulated neurons of the ipsilateral hemisphere. However, adapter tones with positive ILDs, i.e. located on the ipsilateral side, sufficiently excite the neuron to activate the GABA circuit (increasing value of *p_ω_* in membrane Eq. 4). Consequently, such adapter tones decrease the effectiveness of the inputs (by modulating the input weight kernels 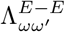 and 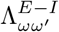 of the excitatory (*E − E*) and inhibitory (*E − I*) inputs, respectively, see Eq. 2 and 3) which leads to a response shift. In other words, the neuron’s response curve and consequently the region with the steepest slope, thus highest sensitivity, shifts towards the adapter tone. This leads to a higher sensitivity for consecutive sounds of the same ILD.

Neural dynamics together with their thresholds and retrograde inhibition are not the only factors that can determine the neuron’s response behavior. Also stimulus latencies play a crucial role [17, 40] as shown in the next experiment.

### Timing

A binaural stimulus simultaneously creates a time and level difference of its sound signal at the sensory level where the relationship between those two cues is monotone (considering broad-band stimuli): Increasing the ITD of a stimulus monotonically decreases the ILD of this stimulus and vice versa. For example, a sound source located at the ipsilateral side of the head produces not only a higher sound level but also a delay of signal arrival time at the contralateral ear. The nature of ILD processing, that is the timed interaction of excitatory and inhibitory inputs, already leads to an essential sensitivity to the arrival time of the inputs. Neurophysiological studies support this in showing that neurons in the LSO are not merely responsive to the level difference but are also sensitive to differences of arrival time of the excitatory and inhibitory inputs [16, 17, 40, 41]. This phenomenon is called *time-intensity trading* of LSO neurons [17, 31]. It describes that LSO neurons can trade the timing of an input with its intensity, thus exhibiting responsiveness to ITDs of incoming signals.

To test to what extend the present model responds to stimuli with various ITDs, how this leads to adaptation and to compare its adaptation effectiveness to the previous adaptation mechanisms, experiment 4 has been designed. In this, we investigate the influence of signal arrival time on model neurons response characteristics.

Network response of a model neuron for stimuli with varying ILDs over ITDs is depicted in Fig. 4**D**. A negative ITD refers to a sound source located on the ipsilateral side of the head, i.e. it arrives at the ipsilateral side first and therefore leads to a delay of the contralateral signal. The effectiveness of an inhibitory input on an excitatory input is decreased for larger time delays between the two input signals, i.e. the excitatory input might be already processed before the other arrives and vice versa. In concert with neurophysiological results, the response of the model LSO neuron strongly depends on the ITD value of the incoming signal (see [16] Fig. 3 for comparison). For ITDs around 600 μs the delayed arrival of the inhibitory input signal causes a reduction of its effectiveness on the response rate. Only inhibitory signals with high level intensity generate a noticeable effect. In contrast, for ITDs around 200 μs the inhibitory signal arrives even before the excitatory signal and leads to a-priori hyperpolarization of the membrane potential. Thus, already low level inhibitory signals cause a reduction of the response rate and lead to a small shift towards positive ILDs. This results in a U-shaped response curve for positive ITD values and a V-shaped response curve for negative ITD values as reported in [17]. As natural stimuli with negative ITDs essentially exhibit positive ILDs and vice versa, the shift in the response rate indicates an adaptation towards more relevant input ranges. In other words, the interaural time difference affects the sensitivity of the neuron for ILDs by means of changing the inhibitory input effectiveness, thus the neuron exhibits time-intensity trading [17, 31]. This effect is generated by an intrinsic adaptation mechanism caused by the statistics of the input stimuli and the neuronal transduction properties rather than via a dedicated mechanism of neural computation.

In order to investigate the effect of this adaptation, Fig. 4**E** depicts the response curves of a simulated LSO neuron for stimuli with various ITD values over ILD. The outer boundaries of the filled area indicate model responses to ITDs (−600 μs and 200 μs) for which the response curve is shifted maximally and thus delimits maximal adaptation. For higher or lower ITDs the response curve is shifted between those maximal values. The shift of the response is maximal 2*dB* for stimuli located at the far ipsilateral side and maximal 38*dB* for stimuli on the far contralateral side. Shifting the response curve facilitates a higher sensitivity to changes in the ILD of the input signal and indicates an adaptation to the perceived stimulus. The limited shift towards positive ILDs results from maximal effectiveness of the inhibitory input. That is, the inhibitory input leads to a maximal strength of the hyperpolarization for only a specific ITD (in this case 200 μs). For stimuli with higher or lower ITDs the hyperpolarization effect vanishes. The larger extend of the rightward shift of the response curve results from a decreasing effectiveness of the inhibitory input. This leads to a lowering of the activation threshold of the neuron. The extend of the rightward shift depends on the intrinsic model parameters (compare Fig. 2**C** and Fig. 2**D**). For an ITD of 600 μs the inhibitory input signal arrives too late, no longer having a noticeable effect on the response. Hence, the neuron’s response curve solely depends on the level intensity of the excitatory input signal (see Fig. 4**F** for a detailed relation between ITD and ILD responsiveness of LSO neurons).

The fact that loud signals are transmitted faster, i.e. they exhibit shorter transduction delays, supports the shift of the response curve towards a more relevant range [17]. Since inputs to model LSO neurons are modeled without considering the transduction of auditory nerve fibres [42] or other lower areas of processing and information about the relative shift of arrival time at the LSO for excitatory and inhibitory inputs is missing, we replicated the experiment of [17] to determine such a latency of inputs. To achieve similar results as in their experiments, level dependent changes of stimulus latency has been set to 10 μs*/dB* (see supplementary Fig. S3). Therefore, we model the mechanism of transduction delays by advancing the onset of stimuli by 10 μs*/dB* for the current experiment. Together with the increased membrane dynamics for high intensity inputs [22] and the influence of ITD values on ILD encoding the response curve of a model LSO neuron is shifted by 38*dB* for inputs with ITD values of −600 μs which leads to a time-intensity trading value of 15 μs*/dB*. Such time-intensity trading is not considered in the previously presented experiments since for these experiments we assume broadband long duration stimuli for which time-intensity trading can be neglected [43].

## Discussion

We introduced a conductance-based model simulating functions of LSO and MNTB neurons for the encoding of ILD values. The model incorporates a novel synaptic modulation mechanism that facilitates dynamic adaptation to stimulus statistics.

In Experiment 1, we demonstrated the effect of model parameters *γ_r_* and *κ_r_* on the response behavior of a model neuron. Specifically we showed that the subtractive inhibition parameter *γ_r_* controls the response range of the neuron whereas the shunting inhibition parameter *κ_r_* determines the slope of the response. These findings indicate that the neuron’s membrane dynamics need to comprise a balanced combination of subtractive and shunting inhibition in order to cover a wide range of input ILDs while simultaneously maintaining a high sensitivity (see Fig. S1). This is important for the adaptation of LSO neurons since it demonstrates that the response curve of neurons can be controlled by the way the inhibitory input is integrated. Therefore, we hypothesise that the retrograde GABA circuit might have different leverages on the subtractive and divisive integration process of a neuron, respectively.

The ratio between the effectiveness of GABAergic inhibition on the excitatory and inhibitory inputs, respectively, is a crucial factor to ensure high coding precision of neurons as demonstrated in Experiment 2. Only a higher effectiveness on the excitatory input than on the inhibitory input leads to a response shift towards more relevant input range. As shown in Fig. 3**D** this ratio strongly depends on model parameters *κ_r_* and *γ_r_* and is therefore depending on whether the inhibitory inputs are shunting or subtractive. To achieve high coding precision despite having small values of *κ_r_* parameter, the ratio between excitatory and inhibitory GABA effectiveness ought to be rather small (1 : 0.3). However, for high *κ_r_* parameter values, the ratio increases. The *γ_r_* parameter values seem to have a minor influence.

Based on the results presented in Fig. 3**A**, we propose that an inverse ratio of the GABA receptor effectiveness, that is *λ_E_* < *λ_I_*, on the excitatory and inhibitory inputs, respectively, can lead to an extension of the neuron’s response range. Simultaneously, it reduces its sensitivity based on the coding precision index (derivative of the neuron’s response curve, high values indicating high sensitivity of the neuron to changes in ILD values, see section GABA parameter strength). Thus, we predict that such a ratio enables a single neuron to tune its response for a wide range of input stimuli. However, adaptation always comes with a trade-off. In this case, the trade-off is between the ability to respond to a wide range of stimuli and the achieved coding precision.

The two presented adaptation abilities of LSO neurons, namely synaptic GABA signaling and sensitivity to ITD values (time-intensity trading), are very different in their underlying principles, however, yield the same effect: adaptation of the response based on stimulus statistics. To quantitatively assess the quality of the adaptation process, we introduced the coding precision value. Based on this index we can now compare the two adaptation processes based on their sensitivity gain and the extend of the response shift (see Fig. 5).

**Fig 5.**
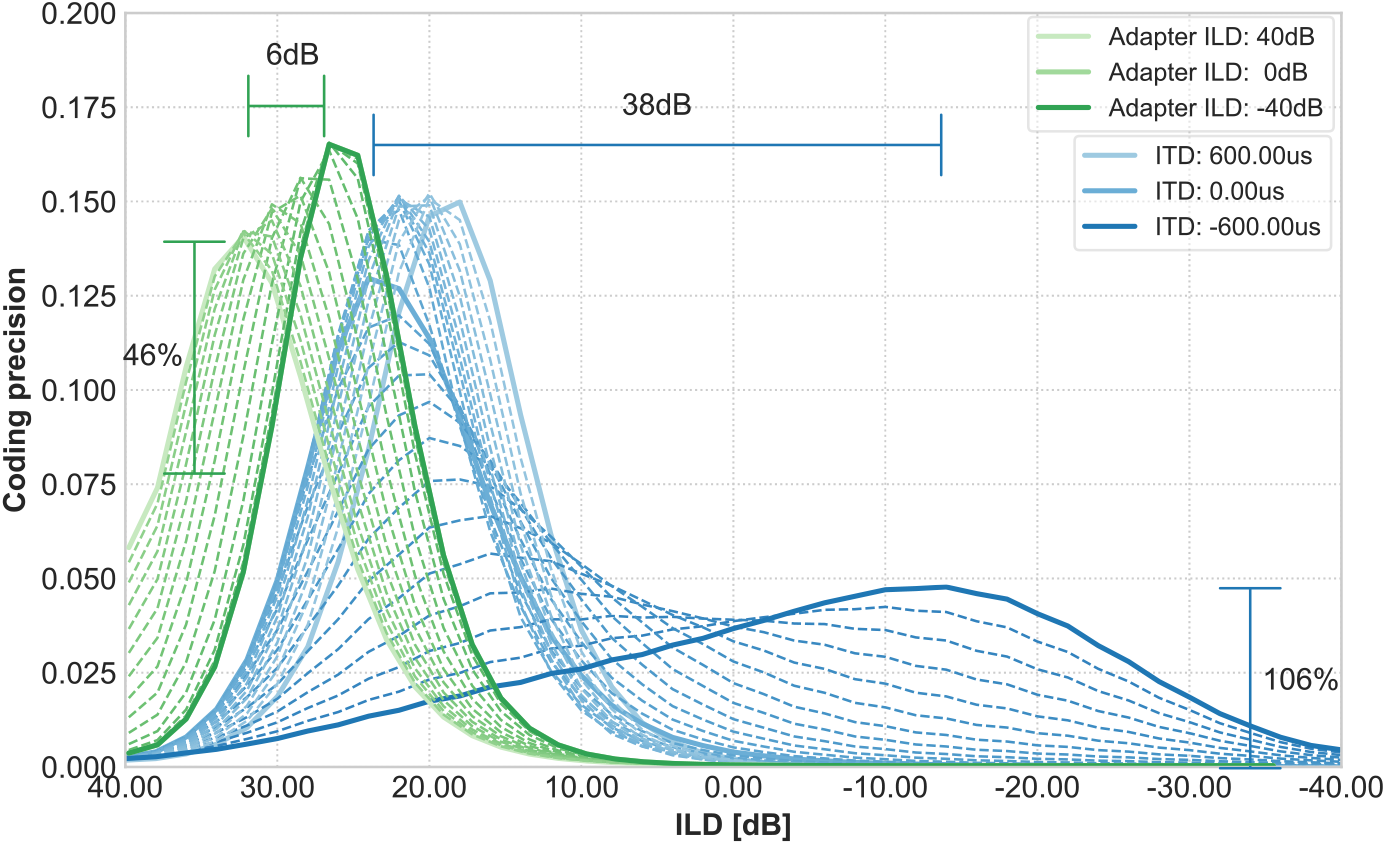
Comparison of coding precision between GABA and ITD induced adaptation. Green lines indicate the coding precision of the adapter tone experiment (no. 3). The peak of the coding precision curve is shifted by a maximum of 6*dB* for increased adapter tone intensity and gains an sensitivity increment of 46%. Blue lines depict the coding precision of the timing experiment (no. 4). The peak of the coding precision curve is shifted by a maximum of 38*dB* between stimuli of different ITDs. The increment of sensitivity for this shift is 106% but is dramatically decreased.

The adaptation induced by the retrograde GABA circuit leads to a shift of 6*dB* of the coding precision maximum towards the ILD of the adapter tone, which corresponds to a gain in sensitivity for ILDs close to the adapter. Particularly, the amplitude of the coding precision and consequently the slope of the response curve at these ILDs is increased by at least 46% of its original value, i.e. when no adaptation takes place. The presented coding precision values are similar to the ones measured in gerbils (Fig. 8 in [20]) when presented with a similar adapter-stimulus combination. In contrast, the adaptation induced by time-intensity trading of LSO neurons (Experiment 4, blue lines) indicates a shift of 38*dB* towards the contralateral side and gains 106% of its original sensitivity, which was basically 0 before.

In a recent study [19], the authors extended a commonly used monaural stimulus paradigm to investigate dynamic range adaptation by a binaural version to probe the effect of such a stimulus in the LSO. Since the applied monaural stimulus has been previously used to investigate monaural adaptation of auditory nerve fibres [44–46] the authors could determine the monaural contribution to the observed adaptation in the LSO. Thereby, derive the LSO specific adaptation capability. The results reported in [19] suggest that the gain control mechanism of LSO neurons spreads the activity over a a population of neurons during adaptation and therefore enhancing the coding efficiency and not necessarily the coding precision. Since the stimulus paradigm applied in their study differed from the one in our adaptation experiment (experiment 3), we can only partly compare the results. A major factor, that we believe contributes to the differing results, is the applied ILD which is in 80% of the cases close to 20*dB* in their study but has a constant value of 40*dB* in our experiment. Thus, the activation of the retrograde GABA signaling circuit is considerably stronger and leads to a higher suppression of consecutive stimuli. When an ILD of 20*dB* is applied in our experiment the results look qualitatively similar to the ones presented in [19] (see supplementary Fig. S4). Furthermore, we demonstrated that the ratio of GABA effectiveness on the excitatory and inhibitory inputs can greatly influence the adaptation ability and the coding precision of a neuron. This variance in GABA ratios between neurons in an LSO population might contribute to the different results.

The presented response shifts can be explained by a reduction or enhancement of the effectiveness of the excitatory and inhibitory inputs depending on the underlying adaptation processes. The reduction is achieved explicitly by inhibiting the excitatory and inhibitory inputs through dynamic temporal adaptation of the weight kernels 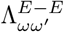 and 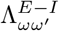. As a consequence, the operational range of the membrane potential (maximal value of *r_ω_*) is reduced which leads to a decreased activation (*g_r_*(*r_ω_*)) of the neuron. In other words, for the same stimulus ILD the neuron exhibits a lower activation level, hence the response over ILD is shifted. In contrast, the enhancement is caused implicitly by a temporal delay of the inhibitory input signal which leads to a reduced influence of this signal in the integration process with the excitatory input signal.

The difference in the maximum of the coding precision function between the adapted and the non-adapted condition is due to the dynamics of the model neuron. If no adaptation takes place the saturation of the membrane potential of the neuron is reached for an ILD of 40*dB*. However, if the GABA circuit attenuates the effectiveness of the inputs, the saturation point of the membrane potential is never reached. Hence, the neuron’s membrane potential is constantly below the saturation point for all input ILDs and can not reach it’s default activation level. In the case of temporally delayed input signals, the inhibitory input exerts less influence which has a similar effect as increasing parameter values *κ_r_* and *γ_r_* (parameters to define the strength of the inhibitory influence). As can be seen in Fig. 2**C** and Fig. 2**D** the change of the inhibitory influence strength changes the slope of the response function. A decreased slope in the ILD response functions leads to a lower amplitude of the coding precision function. Together, the sensitivity to ITD and the retrograde GABA signaling circuit enable a flexible encoding of auditory space and extend the input range of a neuron significantly (6*dB* + 38*dB* = 44*dB*).

The presented adaptation mechanism of retrograde GABA signaling on the example of LSO neurons is very simplistic, thus, is likely to take place in other domains of signal integration as well. For example, neurons in the medial superior olive complex integrate ipsi- and contralateral signals for ITD encoding and could profit from such an adaptation mechanism. Short term synaptic plasticity is a commonly found property of neurons in various brain areas [47, 48]. The introduced mechanism creates a canonical principal for synaptic adaptation and therefore could account for various adaptation mechanisms in the brain [49, 50].

The first single cell model of LSO neurons was introduced by [51] and further investigated in [52]. The authors introduced a detailed multi-compartment model of a LSO neuron which explains its spiking behavior in particular its chopper response at stimulus onset. This response behavior is of minor interest in the present study since we focused on the adaptation mechanisms of LSO neurons and the consequent implications on improved encoding of auditory space which was not the target of Zacksenhouse and colleagues in their studies. For a comprehensive review on other models of the LSO, see [53].

There are several properties of the auditory pathway that are currently not considered in the design of this model, since we believe that these properties do not weaken its predictive power. Nevertheless, we will discuss several of them below that could improve the model and might be targeted in further investigations.

The inputs to the model originate from spherical and globular bushy cells in the ipsi- and contralateral cochlear nucleus, respectively. In our model, we assumed that the characteristics of these inputs are identical, both following the output of auditory nerve fibers of the ipsi- and contralateral side, respectively. That is, the activation changes linearly with the log-intensity of the input [23]. However, this might not necessarily be the case. Spherical bushy cells exhibit a strong response at the onset of a stimulus before settling on a lower response rate for the rest of the stimulus duration (“spike-frequency adaptation”) [33]. In contrast, globular bushy cells’ responses are almost identical to the auditory nerve input [54].

In this paper, we demonstrated how the sensitivity and range of ILD encoding in LSO neurons can be explained by model parameters. Particularly, how different inhibition types (subtractive and shunting) contribute to the response behavior of a model neuron. The neuron’s responsiveness to temporal delays of its inputs indicates an inherent adaptation process which maintains high sensitivity to incoming stimuli. Together with the retrograde GABA signaling circuit, these adaptation mechanisms contribute to the high accuracy of ILD encoding in LSO neurons over a wide range of input stimuli. We compared the coding precision of the two adaptation mechanisms quantitatively and showed that an interplay between the two jointly sensitizes the circuit to enable a response shift towards contra-lateral and ipsi-lateral stimuli, respectively. Thereby a single model neuron can cover almost an entire hemisphere of auditory space. In addition, it saves computational resources by reusing the same population of LSO neurons for various ILD ranges. Most significantly, we proposed an optimal ratio between GABA effectiveness on the excitatory and inhibitory inputs, respectively.

## Supporting information

**S1 Text. ILD Computation.**

**S2 Text Adaptation to lower adapter tone level.**

**S1 Fig. Parameter influence on coding precision.**

**S2 Fig. Parameter sweep for GABA ratio.**

**S3 Fig. Time-intensity trading experiment Park et al. 1996.**

**S4 Fig. Parameter influence on coding precision.**

