## Supplementary Figure 1 for "Computational principles of neural adaptation for binaural signal integration"

**S1 Figure. Parameter influence on coding precision.**

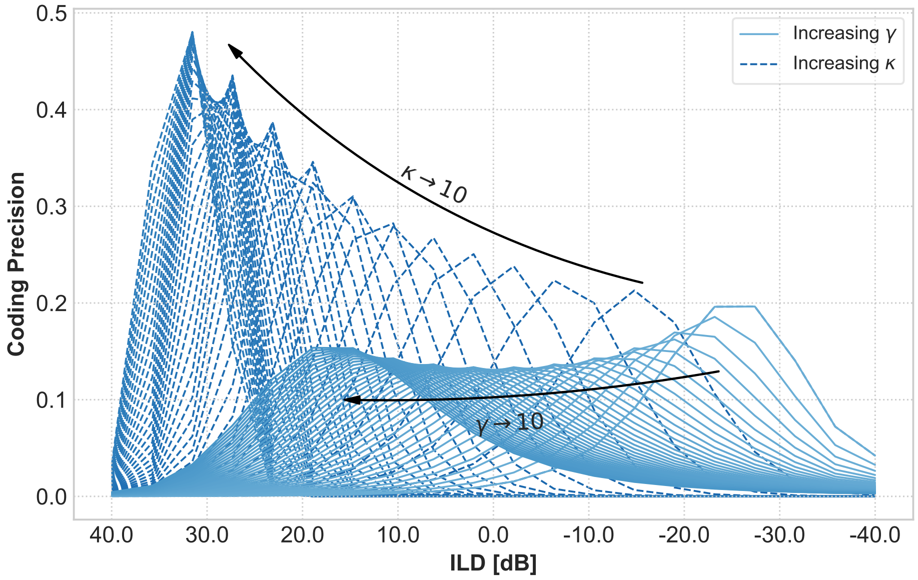


**Parameter influence on coding precision.** Dashed blue lines indicate the coding precision of the model response for changing $\kappa_r$ parameter. For increasing parameter value the coding precision increases while simultaneously the response range decreases (the ILDs that the model exhibits a response). Solid blue lines indicate the coding precision of the model response for changing $\gamma_r$ parameter. For increasing parameter value the coding precision and extensions of the response range remains almost constant. However the response range itself shifts from negative to positive ILD values. A proper ratio of the two parameters facilitates a balance between high coding precision and response range.
