## Supplementary Text 1 for "Computational principles of neural adaptation for binaural signal integration"

**S1 Text. ILD Computation.** For calculating the ILD values of real world stimuli we use the text book definition (e.g [1]) of ILD computation:

$$ILD = 10 * log_{10} \frac{\int_{t=0}^{\infty} s_l^2(t)}{\int_{t=0}^{\infty} s_r^2(t)} \tag{1}$$

where $s_l$ is the signal received at the left ear and $s_r$ is the signal received and the right ear at time $t$.
