## Supplementary Figure 3 for "Computational principles of neural adaptation for binaural signal integration"

**S3 Figure. Time-intensity trading experiment Park et al. 1996.**

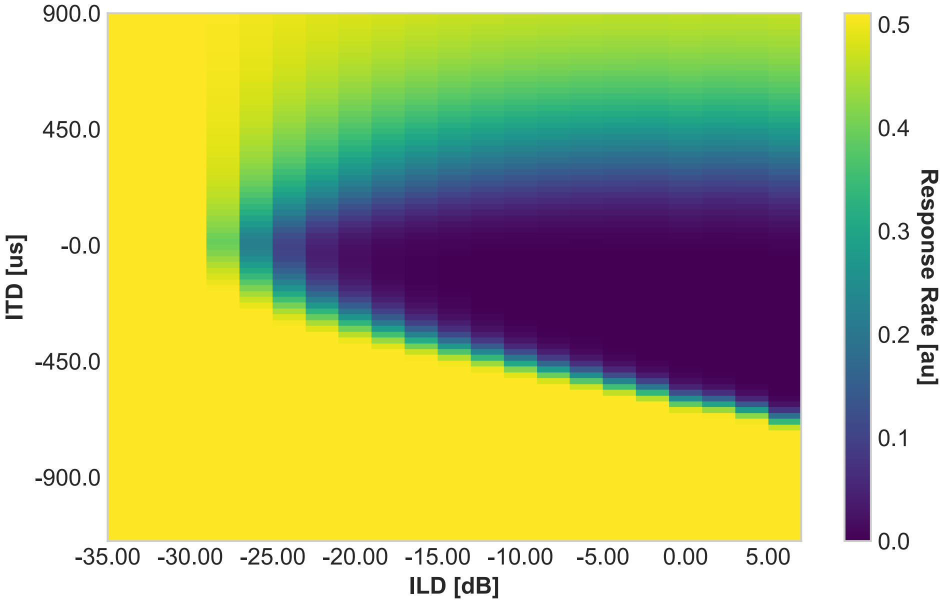


**Time-intensity trading of LSO neurons.** Figure shows simulation result for time-intensity trading experiment similar to [1]. To achieve similar results the model parameters needed to be changed to $\gamma = 10$ and $\kappa = 0$. This change of parameters is reasonable, since the effectiveness of the inhibitory inputs might differ from neuron to neuron [2, 3]. The ipsilateral stimulus intensity was fixed to $35dB$, whereas the contralateral input intensity varied between $0dB$ and $40dB$ thus creating similar ILD values as in the experiment of [1]. The duration of a single stimulus was set to 300 µs. To achieve qualitatively similar results as the authors (compare their Fig. 3) we choose the time-intensity value of the arrival times of the model inputs to be 10 µs/dB. We use this value for the timing experiment (no. 4) to adapt the arrival time of inputs.
