## Supplementary Text 2 for "Computational principles of neural adaptation for binaural signal integration"

### S2 Text Additional experiments

**Point of Complete Inhibition** Park and colleagues [1] investigated properties of neurons in the LSO complex that facilitate the wide variability of ILD responses. The authors presented stimuli of different intensities to mustache bast and measured the response of individual LSO neurons. They reported that a reliable factor to compare the response behavior between neurons is the ILD of *complete inhibition* (PCI) which remains constant for a single LSO neuron but varies significantly from cell to cell (see [1] Fig. 3). That is, the slope and maximum saturation value might differ depending on the presented stimuli level, whereas the point for which the LSO neuron is completely deactivated remains constant over different input intensities. Therefore, it can serve as a unique index to describe the neuron's response characteristic. To compare their findings with results of the present model, figure 1 shows the model response to similar stimuli. That is, a signal of constant level intensity is presented on the ipsilateral side (as noted in the legend) and a signal of consecutively increasing level intensity is presented on the contralateral side. The response of a model neuron varies with varying input intensities but the point of complete inhibition remains constant. This result also confirms the threshold hypothesis of Reed and Blum [2], which states that the difference of thresholds for excitatory and inhibitory input corresponds with the point of complete inhibition. That is, the effectiveness of the excitatory and inhibitory inputs on a neuron's dynamics determine its point of complete inhibition.

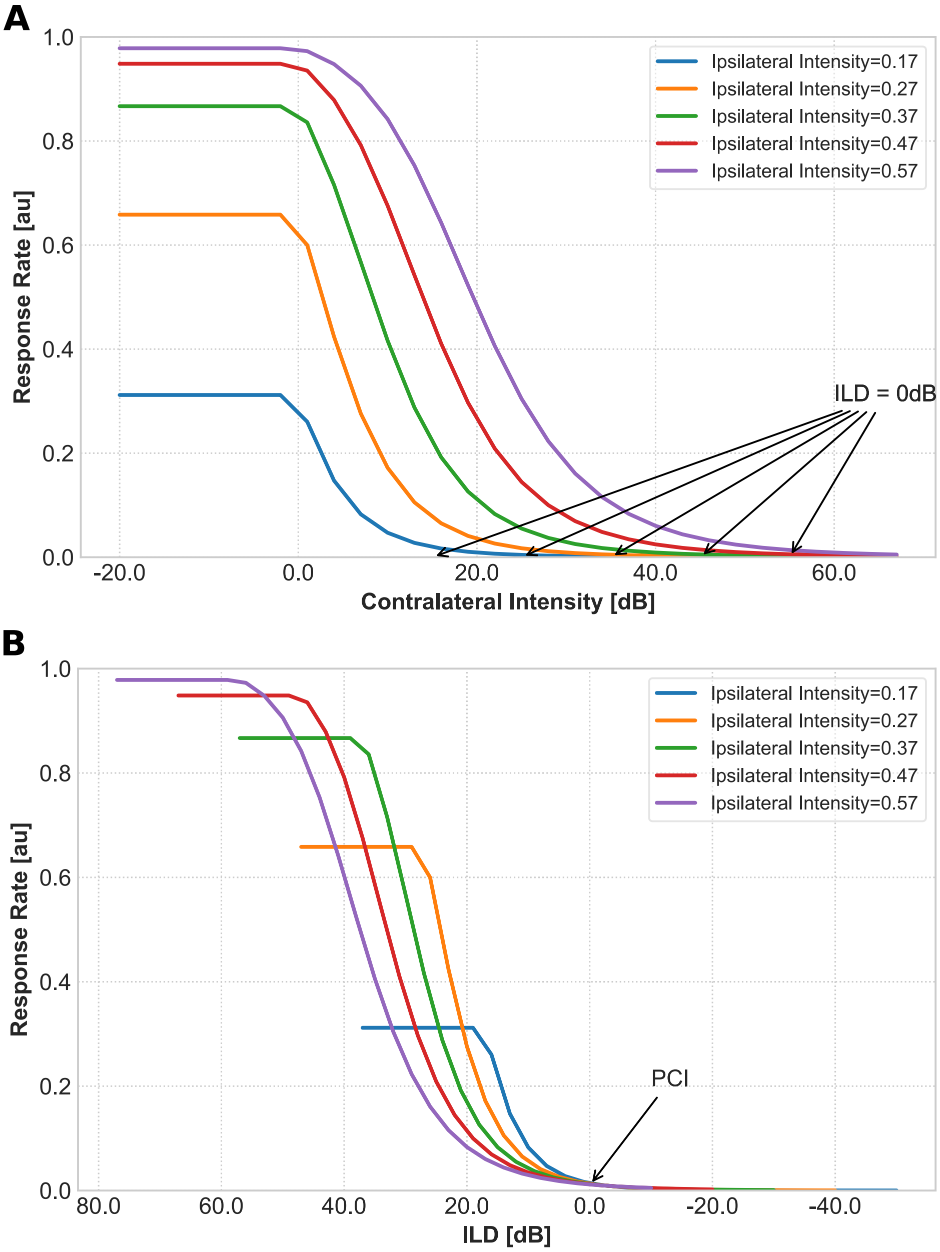


Figure 1: **Point of complete inhibition** (A) Neuron response for different ipsilateral input intensities over contralateral input intensities as in [1] their Fig. 3. (B) Summary of results in the standard ILD plot. Different colors indicate various ipsilateral input intensities.
