## Supplementary Figure 4 for "Computational principles of neural adaptation for binaural signal integration"

**S4 Figure. Adaptation to lower adapter tone levels.**

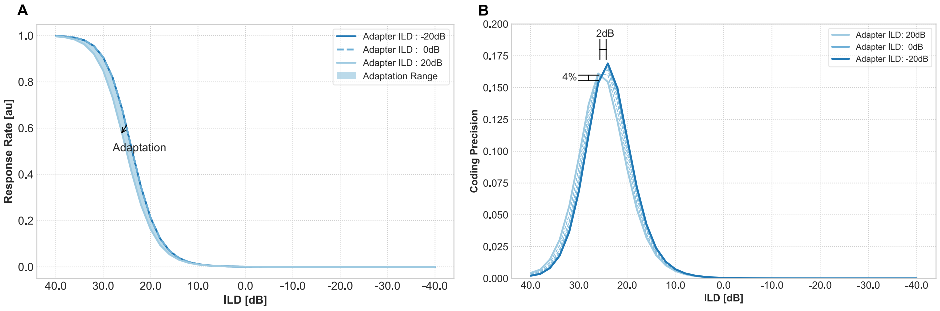


**Adaptation to lower adapter tone levels.** Same stimulus paradigm as in experiment no. 3 (Fig. 4**B**) except that the adaptor tone intensity was limited to be in range $-20dB$ and $+20dB$, which is similar to the highest ILD value used in [1]. (A) the maximum adaptation range decreases and so does the coding precision value (B) which creates qualitatively similar results to then one Gleiss and colleagues found ([1] compare their Fig. 4).
