## Supplementary Figure 2 for "Computational principles of neural adaptation for binaural signal integration"

**S2 Figure. Adaptation for plausible ITD values.**

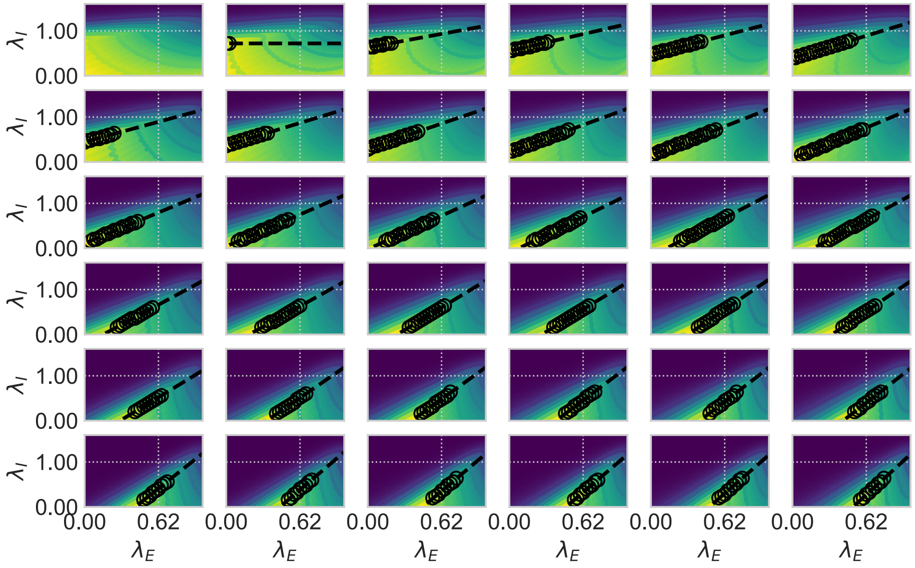


**Parameter influence on GABA ratio** Head map plots for different combinations of model parameter $\kappa_r$ and $\gamma_r$ values are shown. Each plot depicts an exhaustive parameter evaluation of the $GABA$ parameters $\lambda_E, \lambda_I$ . Each point in the heat map indicates the coding precision value for a certain combination of $\lambda_E$ (abscissa) and $\lambda_I$ (ordinate). We assumed that the neuron's task is to achieve a preferably high coding precision value. Black dots depict these values (the maxima of the map). A linear function (dashed line) was fitted to those points (linear regression) and the slope and bias calculated.
